# Failure through expanding voids in bacterial streamers

**DOI:** 10.1101/225367

**Authors:** Ishita Biswas, Ranajay Ghosh, Mohtada Sadrzadeh, Aloke Kumar

## Abstract

We investigate the failure of thick bacterial floc-mediated streamers in a microfluidic device with micro-pillars. We found that streamers could fail due to the growth of voids in the biomass that originate near the pillar walls. The quantification of void growth was made possible by the use of 200 nm fluorescent polystyrene beads. The beads get trapped in the extra-cellular matrix of the streamer biomass and act as tracers. Void growth time-scales could be characterized into short-time scales and long time-scales and the crack/void propagation showed several instances of fracture-arrest ultimately leading to a catastrophic failure of the entire streamer structure. This mode of fracture stands in strong contrast to necking-type instability observed before in streamers.

## Introduction

Bacterial streamers are filamentous biofilm-like structures that are usually known to form under sustained hydrodynamic flows ^1^. Like biofilms, streamers consist of bacterial cells embedded in a matrix of self-secreted extracellular polymeric substances (EPS) and are excellent examples of soft materials of biological origin. Due to their morphology, streamers can colonize closed channels significantly faster than surface-hugging biofilms; recently streamers forming in very low Reynolds number conditions (*Re* ≲ 1) have been implicated for their role in rapid fouling of biomedical devices ^2–4^, filtration units ^5,6^ and even colonization of porous media ^1,7,8^. In many of these applications, a better understanding of deformation, fracture and failure of streamers is crucial ^5^. Valiei et al. ^8^ had reported failure and disintegration of streamers in microfluidic device, but this phenomenon was not discussed in detail. Das and Kumar ^9^ had investigated instabilities and break-up of streamers when they were idealized as highly viscous liquid jets. In a later work, Biswas et al. ^5^ utilized a microfluidic device with micro-pillars to investigate far-from-wall failures of streamers. They focused exclusively on ‘thin’ streamers, i.e. streamers whose aspect ratio (η), i.e. ratio of longitudinal to transverse characteristic length, is typically >10, and found that these streamers could fail through a necking like failure mode in steady flow typical of ductile materials under creep. They also found a power law relationship between the critical strain (strain at instability) and fluid velocity scale, which yielded valuable insights into material behavior. Recently, Hassanpourfard et al. ^10^ showed that failure of streamers is not limited to the earlier stages but can also be found in the final clogged state of the device where localized failures lead to intricate water-channels coursing through the clogged biomass. Despite these studies, it is almost certain that more failure modes exist. This is because streamers like other biofilms represent a composite and extremely heterogeneous, active soft material. However, reporting and quantification of these phenomena is sparse.

One of the most fundamental challenges stems from the visualization issues since the EPS matrix embedding the microbe is very difficult to image due to its transparent nature. Furthermore, the nature of this type of set up results in several overlapping sources of nonlinearity in addition to failure and instability. These include creep behavior of the polymeric EPS, fluid-structure interaction, moving interfaces and life processes, all of which cannot be independently controlled easily. In addition, time scale of streamer formation can vary considerably ^3,7,8,11^ and very long-time scales can let significant changes in the background conditions affecting these nonlinear behaviors.

In this communication we report an entirely new type of streamer failure mechanism not observed before. This type of failure originates near the micropillar wall, rather than further downstream as previously noted, and yet distinct from shear failure at the micro pillar wall typically seen at higher flow rates ^5^. We use floc-mediated ^5,7,10^ rather than biofilm mediated ^3,8^ streamer formation, where either of these refer to the mode of inception of the streamers ^1^. Floc mediation allows for rapid streamer formation which helps us isolate mechanical factors and reduce streamer formation time thus reducing biophysical complications such as cell division ^7^ A microfluidic device was specially fabricated so as to allow streamers to freely form from micro pillars into the downstream flow without any more attaching surfaces on the free side. Our imaging clearly showed that the inception of failure occurs with a pronounced void almost with the geometry of a small coin shaped crack near the point of attachment and only after the streamer structure was already well formed (i.e. when streamer length is several times the pillar characteristic length). Once this ‘crack’ was observed, it was found to rapidly extend resembling crack propagation quickly rupturing the streamer. We found this behavior repeatedly, always originating near the pillars and only for some but not all streamers. This failure mode did not occur anywhere farther down the streamer length. The flow rate was kept constant for individual experiments. To the best of our knowledge, this is first report on this type of failure mode in streamers formed in micro fluidic environments.

## Materials and Methods

### Microfluidic chip fabrication

A PDMS (Polydimethylsiloxane) microfluidic device was made by using traditional photolithography technique for this study. A 4″ silicon wafer was utilized to make the master mold of the microfluidic device. The microfluidic device design consists of a straight channel with a singlet inlet and outlet (Fig. 1a). The length of the channel, *L*, was 11.5 mm and its width, *W*, was 0.436 mm. In the central section of the channel, 14 micro-pillars were arranged in a staggered pattern. Using photolithography process as detailed by Hasanpourdfard et al. ^12^ PDMS microchannels were prepared from a silicon master. The PDMS micro-pillars had a diameter, *d*, of 50 *μ*m height, *h*, of 50 *μ*m and the pore-gap, *p*, was 10 *μ*m (Fig. 1b). The dimensions *w_1_* and *w_2_* demarcated in Fig. 1b were 60 *μ*m and 104 *μ*m respectively. Glass cover-slips (thickness 0.13 to 0.17mm) (Fisher Scientific, ON, Canada) and PDMS were bonded by using oxygen plasma and the devise was then annealed at 70°C for 10 minutes to seal perfectly.

**Figure 1:**
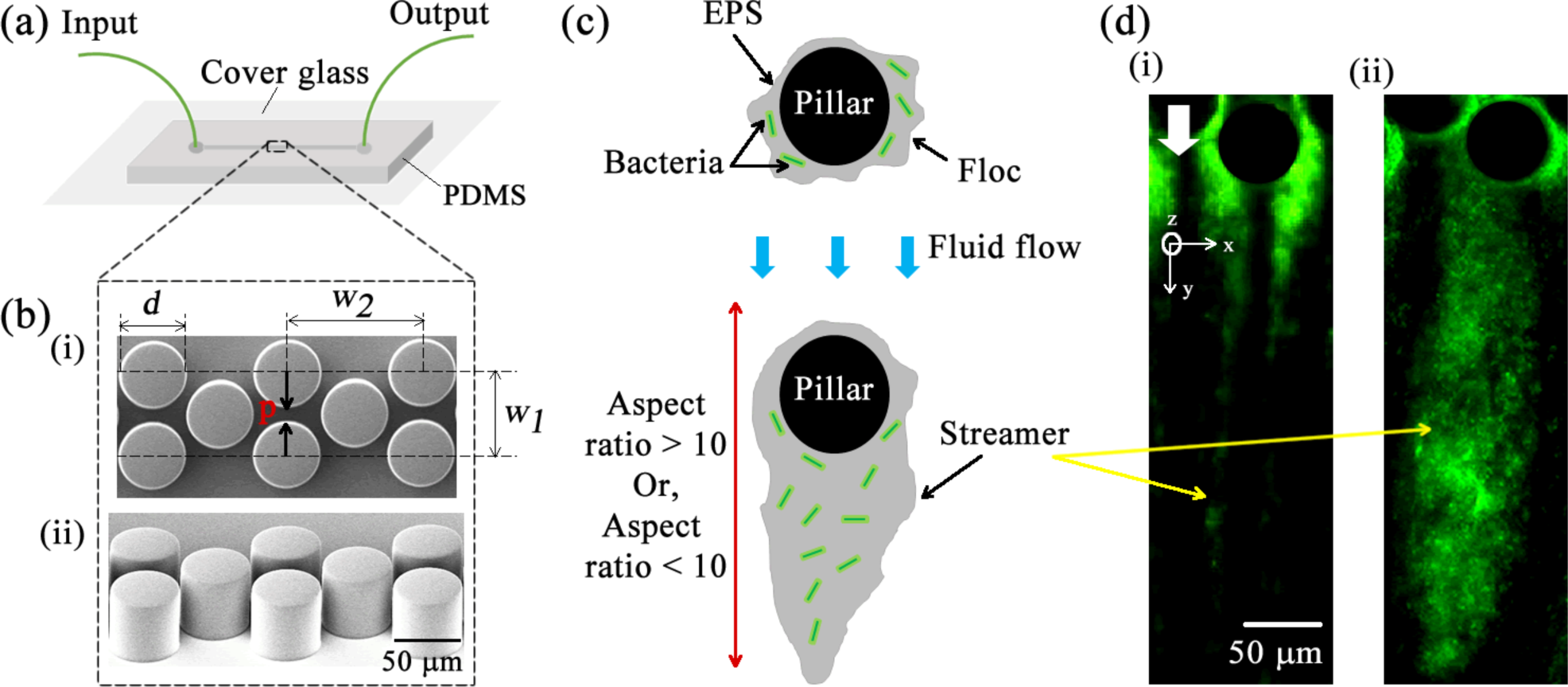
(a) Schematic of microfluidic chip (b) SEM image of the micro pillars. (i) Top-view. The dimensions are *d* = 50 μm, *w*_1_ = 60 μm, *w*_2_ = 104 μm, height of the pillars, *h =* 50 μm and gap between pillars is *p* = 10 μm and (ii) Side-view (c) Schematic of streamer formation. Bacterial flocs attach to the micropillars. Fluid flow generates traction forces that causes the biomass to extrude out in the form of filamentous structures i.e. streamers. In our chip design, one end of the streamer is attached to the pillar wall, while the other end extends downstream. (d) Green fluorescent microscopy images of i) thin streamer, aspect ratio >10 and ii) thick streamer, aspect ratio <10. The downward flow of fluid is indicated by the white arrow.

### Bacteria culture preparation

We used *Pseudomonas fluorescens* CHA0 (wild type) ^13^ bacteria strain for this study. This gramnegative aerobic bacteria is found naturally in water and soil, and plays a vital role in plant health ^14^ The bacteria was genetically modified to produce green fluorescent protein (GFP) constitutively and hence appears green under fluorescence imaging. The bacteria strain was taken from –80°C collection and streaked on Luria Bertani (LB) agar plate in a zigzag pattern. The plate was incubated overnight at 30°C. Subsequently, a single colony was taken from the plate and put into liquid LB broth and incubated it for 26 hours in a shaking incubator (Fisher Scientific, ON, Canada) at 150 rpm and 30°C. The longer incubation time was employed so that bacterial flocs formed ^7^. This bacteria culture was mixed with 200 nm red fluorescent amine-coated polystyrene (PS) microspheres (Thermo Fisher Scientific, MA, USA). The final concentration of PS beads in the mixture was 0.02% (v/v) and this mixture was injected into the microfluidic chip.

### Microscopy

The bacterial solution was injected to the microfluidic chip by using a syringe pump (Harvard Apparatus, ON, Canada). The microfluidic chip was placed under an inverted fluorescence microscopy system (Nikon Eclipse Ti) to observe the streamer behavior during the experiment. Confocal images were captured using an inverted spinning-disk confocal microscope (Olympus IX83). For epi-fluorescence imaging GFP Long-pass green filter cube (Nikon and Olympus) and Texas red filter cube (Nikon and Olympus) were used. Streamer breaking events were recorded at a rate of 10 frames per second. Image processing was performed by using NIS-Element AR software interface (Nikon). The software is capable of fitting ellipses to images.

## Results

When the *P. fluorescens* bacteria solution was injected through the microfluidic channel, streamers form downstream of the micro-pillars with a characteristic streamer formation time scale, *t_form_*. For a floc laden bacteria solution, as in this experiment, *t_form_*∼*O*(10^−1^ – 10^0^ *s*). Streamer formation in these cases occurs because the flocs adhere to the micro-pillar walls and are rapidly sheared into the form of streamers by traction forces generated by the flow (Fig. 1c). This mode of streamer formation has been investigated and characterized previously ^5,7^ Figure 1d shows two different streamers formed in the device, when the bacteria solution was injected through the chip a constant volumetric flow rate (*Q*) by using a syringe pump. The corresponding global velocity scale, 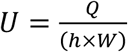, and Reynolds number, *Re*, were 3.75 × 10^-3^ m/s and ∼10^-3^ respectively. The two streamers differ in their morphology. The streamer marked ‘(i)’ is an example of thin streamers, whose aspect ratio (ratio of longitudinal to transverse length) η > 10. The longitudinal and transverse length of a streamer is measured along the y-axis and x-axis respectively as shown in Fig. 1d. The streamer marked ‘(ii)’ is an example of a ‘thick’ streamer, η < 10. A constant flow rate is maintained for the entire duration of an experiment. After the streamers form, a quiescent period, *t*_o_, is usually observed, where no significant deformation of streamers is observed. At the end of this time period, *t*_o_∼*O*(10^2^ – 10^3^ *s*), subsequently noticeable deformation in the streamer mass can be observed often leading to the disintegration of streamer structure. One mode of failure is the already reported necking type failure prevalent far from the pillar wall ^5^. However, yet another mode of failure was discovered in these experiments which occurred near the micro pillar wall, discussed below. Figure 2a depicts a composite confocal microscopy image of the void in streamer mass. This composite image was created by overlaying the image obtained using the two different filter cubes, and thus both bacteria (green) and PS beads (red) can be seen. As the time resolution of confocal microscopy imaging is comparable to the streamer deformation, Fig. 2a can only yield qualitative insight into the geometry of the void. As we can see the void spans the entirety of the *z*-axis depth of the streamer and thus ‘open’ to the background flow. Figure 2a also shows that the void geometry can be better approximated when imaging using the Texas Red filter cube is performed. The amine-coated PS beads being significantly smaller than the bacterium easily embed themselves in the EPS matrix, thus serving as useful markers ^5,7^. Since confocal microscopy does not allow us to investigate the shape of the void, for the rest of the investigation epi-fluorescence optical microscopy was utilized. In Fig. 2b, we show the moment of onset of a void in the streamer mass after approximately 71 minutes of initiation of experiment. To ensure that the proper void boundary is tracked by image processing algorithms, imaging was performed using the Texas red filter so that the 200 nm PS beads embedded within the EPS matrix are illuminated. Within the limits of optical microscopy and the microscale environment of the microfluidic channel, this allows for the best estimate of the boundary of the growing void.

**Figure 2:**
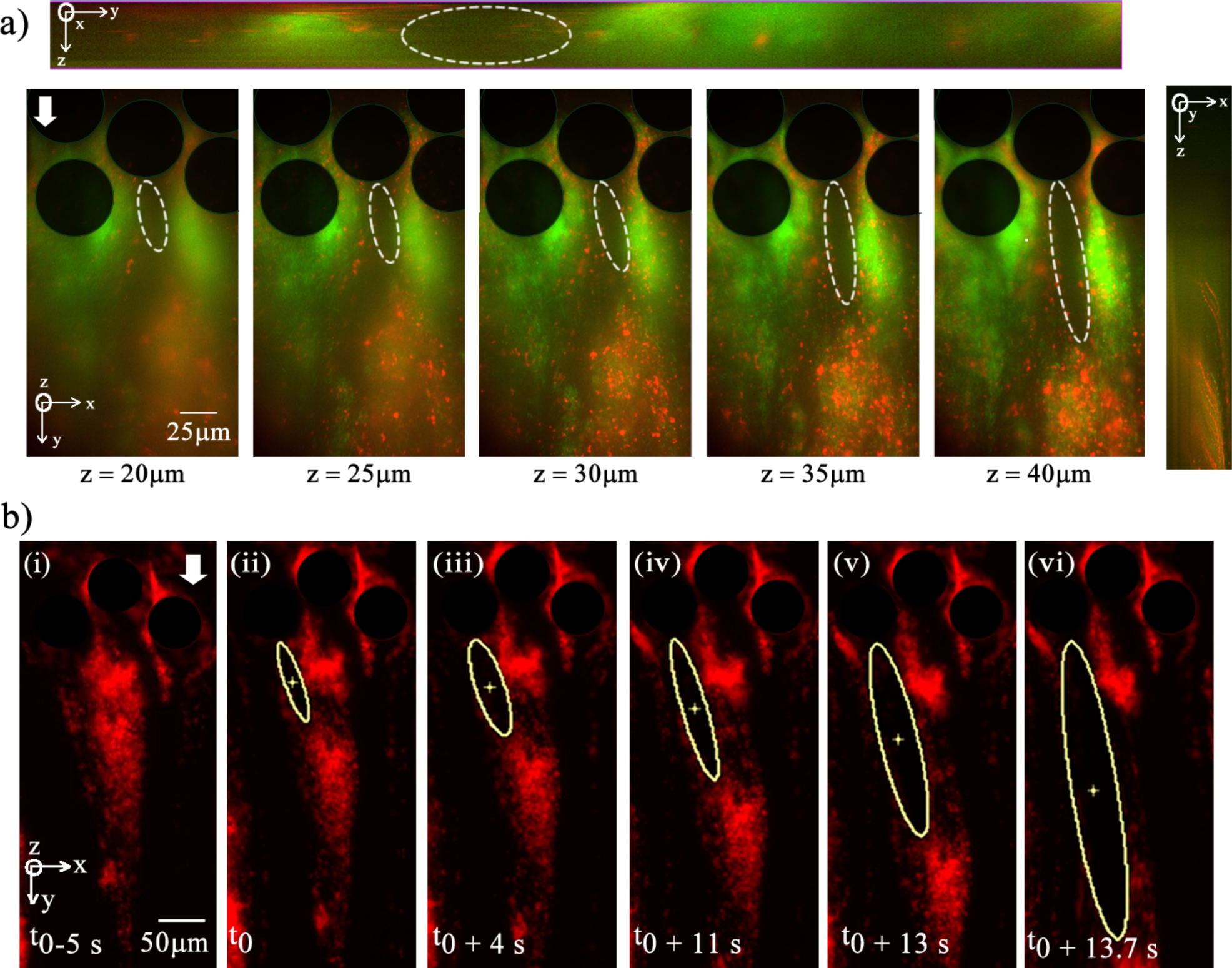
Fracture progression in a streamer a) Composite confocal slices of a void, showing that the void spans the height of the microchannel. Confocal sidebars show the *y-z* plane (top) and *z-x* plane (right). b) At mid-height a growing void is tracked by fitting ellipses. (i-vi) show progression of cavity. The global flow velocity is *U* = 8.92 × 10^-3^ m/s and *t*_0_ = 1 hour 11 minute.

After the onset, the void grows rapidly with time both transversely and longitudinally although the latter being much more pronounced. Figure 2b shows that the void grows with time eventually culminating its progress to the extremities of streamer leading to breakage. To further quantify the breaking phenomena, we tracked the temporal growth the void. Figure 2b also tracks the void and quantifies its deformation by fitting an ellipse to the growing cavity. To quantify the void we define, normalized crack length λ = *a*/*a*_o_ where *a*_o_ is the initial crack length and *a* is the current crack length (both measured along the ellipse’s longitudinal axis).

It is possible to associate with every failure event a fracture time scale (*t_fracture_*). Although, there was a spread in *t_fracture_*, it was possible to bin many of the events in two distinct sets. – one which occurs rather early in the life of streamer (small time-scale fracture-STF) and the other that is observed much later (long time-scale fracture - LTF) (See Fig. S1 for relevant histogram). The STF events that occurred such that *t_fracture_*∼*O*(10^1^ *s*) and their crack propagation dynamics is depicted in Fig 3a. The LTF events occurred such that the set of fracture events occurred slower and for these events *t_fracture_*∼*O*(10^2^ *s*) (Fig. 3b). Fracture time-scales that lie in-between these two extremes were also observed, and similar crack-propagation behavior was observed (data not shown). In addition to crack extension with time, we also plot crack propagation velocity normalized with background flow with time and crack length for both LTF and STF events. This is depicted in Fig. 4.

**Figure 3:**
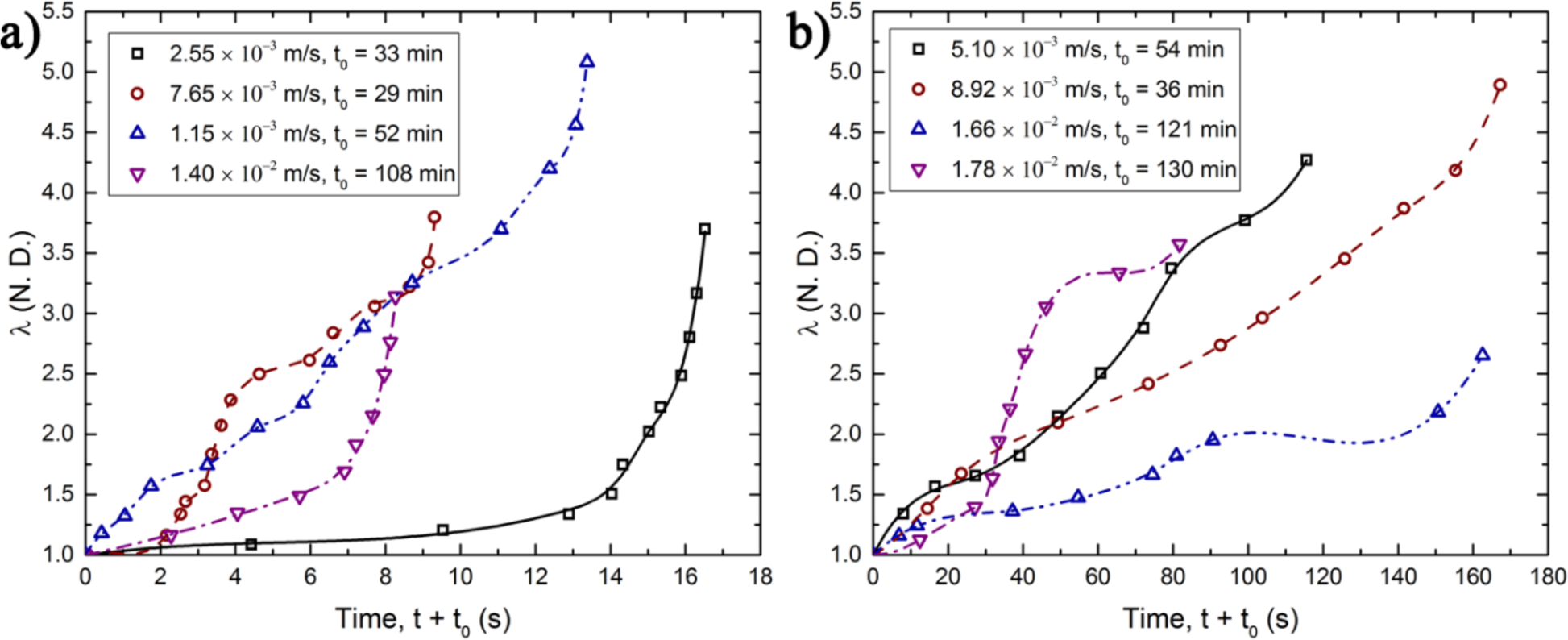
(a) *λ* versus time plot for short time scale fractures. Legend shows the velocity scale (*U*) and *t*_0_ for the different cases. Markers indicate measured data and the lines represent B-spline fits. (c) *λ* versus time plot for long time scale fractures. Legend shows the velocity scale (*U*) and *t*_0_ for the different cases.

**Figure 4:**
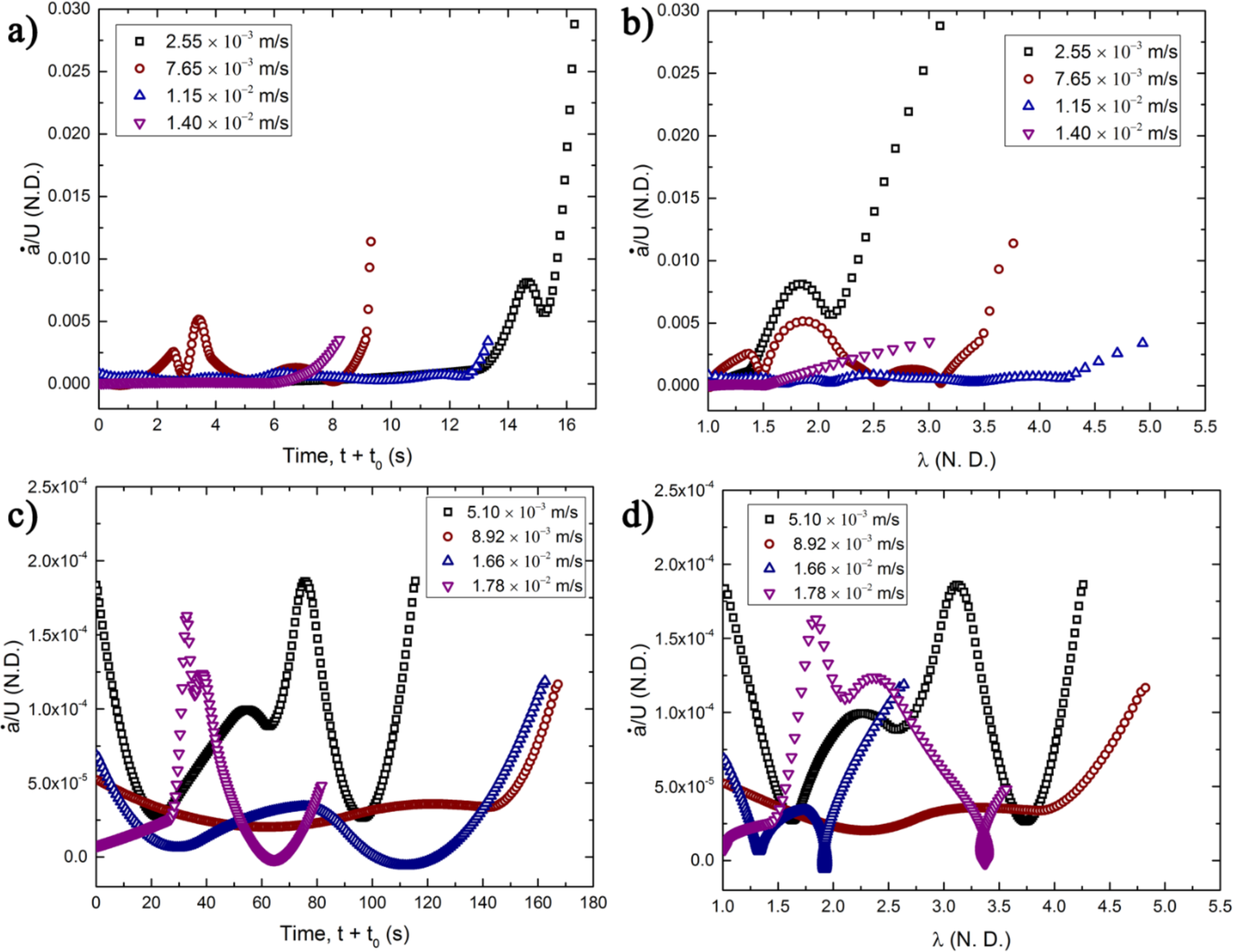
(a) Temporal variation of the normalized crack velocity (*ȧ*/*U*) for STF events. The legend indicates the value of *U* at which the (respective) experiment was conducted. (b) Variation of *ȧ*/*U* versus *λ* for STF events. (c) Temporal variation of the normalized crack velocity *ȧ*/*U* for LTF events. (d) Variation of *ȧ*/*U* versus *λ* for LTF events.

## Discussions

This work highlights a hitherto unreported route of streamer disintegration – one starting right at the base of streamer near the pillar. It is interesting to note that the background fluid flow in the current problem would lead to a traction field in the crack propagation direction, which is not a very favorable crack opening force field. Therefore, by itself it is not the immediate cause of the instability. However, as the void is ‘open’, fluid can enter the void causing the crack propagation to occur. This explains why this form of instability is not found far from the wall since a steady source of fluid ingress is necessary. It also explains why this form of failure does not preclude the possibility of other types of far from wall failure modes since they are entirely independent modes of failure. At the same time, the success of crack propagation would also depend critically on the geometry of the initial void since it not only controls the tip stress conditions which can drive the crack but also the fluid variables inside the initial void. Therefore, the role of geometry of the initial flaw is even more important than traditional fracture propagation experiments. This double sensitivity to imperfections explains the relatively wide scatter in overall crack extension-time and crack velocity-time behavior when flow conditions are changed even slightly even though the resulting traction filed may not be the immediate cause of instability as explained earlier. However, in spite of this difference, the crack propagation characteristics have some broadly similarities. For instance, from Fig. 3(a) and (b), crack extension with time for both STF and LTF show similar nonlinear regimes of propagation for most of the observation– initial linear part, followed by a plateau and thereafter a final catastrophic instability leading to complete disintegration of the streamer. This behavior can be further analyzed using crack propagation velocity plots, Fig. 4a-d, which show an arresting behavior after initial instability. For STF events, crack arrest events are evidenced in both crack length vs time, Fig. 4a as well as crack velocity vs the normalized crack length Fig. 4b. Both of these figures show clearly marked peaks in velocity indicating crack arrest at an intermediate time. For LTF events, the crack arrest events become more frequent as multiple peaks in velocity (Fig. 4c,d) are visible indicating repeated crack arrest after initial propagation. We describe these zones below.

The first zone is the initial crack propagation zone which marks the beginning of the instability and lengthening of the initial flaw. However, as soon as the instability begins likely due to accumulation of damage due to creep at the fracture tip, there is a decrease in crack propagation velocity with time leading to a plateau zone. Note that this plateau is more pronounced for the LTF than the STF. After some time, the plateau zone gives way to a final monotonic increase in crack length for both types of fractures leading to the final disintegration of the streamer. These plots clearly show that as the crack length increases, there is an initial resistance to the crack propagation. This behavior is also seen in many materials which is often a clear indication of nonlinear behavior at the crack tip ^15–17^. This figure also shows that within these broad three-zone behaviors there are further intricacies such as multiple micro regions of crack growth resistance in the plateau zone. However, in all cases, these resistance mechanisms finally fail to stop the crack which then transitions into the final runaway instability zone from which there is no recovery. The streamers ultimately disintegrate.

The broad reason for initial resistance to crack growth likely shares its physical origin to polymeric composite materials which results in overall similar behavior between LTF and STF behavior ^18–20^. However, lack of clear trends in STF may indicate that these streamers may have much higher heterogeneity in their structure leading to more flaws which precipitate the fracture early in their life. This would also explain why the initial crack propagation velocity is much higher for STF, Fig. 3a,b which can be attributed to much weaker interfaces and flaws.

We refrain from developing a complete multi physics model for this system in this communication since its validation would require more precise measurements of flow, material and diffusive parameters.

## Conclusions

In this communication, we discovered a yet unreported failure mode of streamer disintegration which originated near the micro-pillar walls and is yet distinct from simple shear failure or tearing. This failure was observed to co-exist with other previously reported failure modes in streamers. This further supports the notion that this mode of failure is completely independent from previously reported data. We quantify the void/crack extension behavior using PS micro beads as tracers to aid microscopy and conclude that the crack is through the thickness of the streamer and its propagation is marked by very well defined three different zones of propagation. Breaking events occurred at various times after the experiments began but two high frequency zones characterized by a short time scale and another by longer time scales were clearly observed. It is unclear why failure inception is suppressed in the intermediate time scales. The crack/void propagation went through several cycles of propagation and arrest ultimately leading to a catastrophic disintegration/splitting of the entire streamer structure. This mode of streamer disintegration which occurs near the wall can have important implications in spreading of biomass downstream in filtration systems, porous media and biomedical devices.

